# Small-molecule binding to an intrinsically disordered protein revealed by experimental NMR ^19^F transverse spin-relaxation

**DOI:** 10.1101/2023.05.03.539297

**Authors:** Gabriella T. Heller, Vaibhav Kumar Shukla, Angelo M. Figueiredo, D. Flemming Hansen

**Author notes:** Correspondence should be addressed to D.F.H.

## Abstract

Intrinsically disordered proteins are highly dynamic biomolecules that rapidly interconvert between many structural conformations. Traditionally, these proteins have been considered un-druggable because of their lack of classical long-lived binding pockets. Recent evidence suggests that intrinsically disordered proteins can bind small, drug-like molecules, however, there are limited approaches to characterize these interactions experimentally. Here we demonstrate that ligand-detected ^19^F transverse relaxation rates (*R*_2_) obtained from Nuclear Magnetic Resonance spectroscopy are highly sensitive to the interaction between a small-molecule and an intrinsically disordered protein, in contrast to chemical shift perturbations which are minimally sensitive for this interaction. With this method, we show that the small molecule, 5-fluoroindole, interacts with the disordered domains of non-structural protein 5A from hepatitis C virus with a *K*_d_ of 260 ± 110 μM. We also demonstrate that 5-fluoroindole remains highly dynamic in the bound form. Our findings suggest that ligand-detected ^19^F transverse relaxation measurements could represent a highly effective screening strategy to identify molecules capable of interacting with these traditionally elusive, dynamic biomolecules.

## Introduction

Interactions between small-molecules and proteins are ubiquitous in biology, underpinning signaling, metabolism, and therapeutic intervention. Despite the importance of these interactions, the current understanding is largely based on interactions between small molecules and structured proteins, whereby small ligands are often described as binding into well-defined binding pockets. Nevertheless, many proteins, including approximately one-third of human proteins, have long regions which lack such long-lived binding pockets. These include highly dynamic intrinsically disordered proteins (IDPs) that rapidly interconvert between different structures and which are highly abundant in eukaryotes.^[2-4]^ Some IDPs have been shown to interact with small molecules^[5-8]^ thus opening up an enormous class of potential drug targets.^[9]^ The current understanding of the biophysical mechanisms that underpin the interactions between small molecules and IDPs is largely based on theoretical molecular dynamics (MD) simulations that, in turn, have demonstrated that these interactions are highly dynamic.^[5-6, 8, 10-12]^ While offering important insight, MD simulations of IDPs and their interactions, are computationally expensive, require long timescales to reach convergence,^[13-14]^ and suffer from force field inaccuracies.^[13-15]^ Similarly, there is currently a lack of available experimental techniques suitable for generally detecting and characterizing interactions between IDPs and small molecules. Together, the lack of available and accessible approaches forms a major bottleneck to discovering new IDP/small-molecule interactions, uncovering the underlying binding mechanisms, and exploiting these interactions for potential treatment.

## Results and discussion

### ^19^F transverse spin-relaxation is sensitive to binding

To establish sensitive experimental methods to screen for and characterize interactions between small-molecules and IDPs, we utilized the disordered domains 2 and 3 from the non-structural protein 5A (NS5A-D2D3) from the hepatitis C virus (JFH-1 genotype) as a model system^[16-18]^. We focused on solution-state Nuclear Magnetic Resonance (NMR) spectroscopy, since this technique uniquely provides atomic-resolution insight on biomolecular interactions in physiological environments,^[19-20]^ without the need to apply large labels nor localize the molecules on a surface; both of which may alter the structural ensemble and thus behavior of the IDP.

Standard NMR experiments, such as ligand-detected chemical shift perturbations are commonly employed to screen and assess the binding of small-molecules to structured proteins.^[21]^ However, chemical shifts report on the local environment of the nuclei in question which, in turn, are averaged over time and over all the molecules in solution. Given the proposed dynamic nature of the interactions between IDPs and small-molecule ligands,^[5-6, 8, 11]^ we rationalized that NMR parameters that report on ligand dynamics and exchange might be more sensitive to detect IDP/small-molecule binding than chemical shifts.^[22]^

We chose to assess the binding of ^19^F-containing small molecules to NS5A-D2D3 by initially quantifying ^19^F NMR transverse spin-relaxation rates *R*_2,eff_ via a CPMG-based *R*_2_ experiment^[1, 23-24]^. Using ^19^F instead of ^1^H as a probe has the advantage that there are no background signals in the NMR spectra, from buffer components nor protein, and these spectra therefore exclusively report on the small molecule in question. Using this approach, we identified that 5-fluoroindole interacts with NS5A-D2D3 (**Figure 1)**. Specifically, as the concentration of NS5A-D2D3 is increased in samples containing 50 μM 5-fluoroindole, the effective transverse relaxation rate of the ^19^F nucleus in 5-fluoroin-dole, *R*_2,eff_, increases systematically (**Figure 1a-c)**, demonstrating that ^19^F transverse relaxation rates allow for a sensitive detection of small-molecule binding to IDPs.

**Figure 1.**
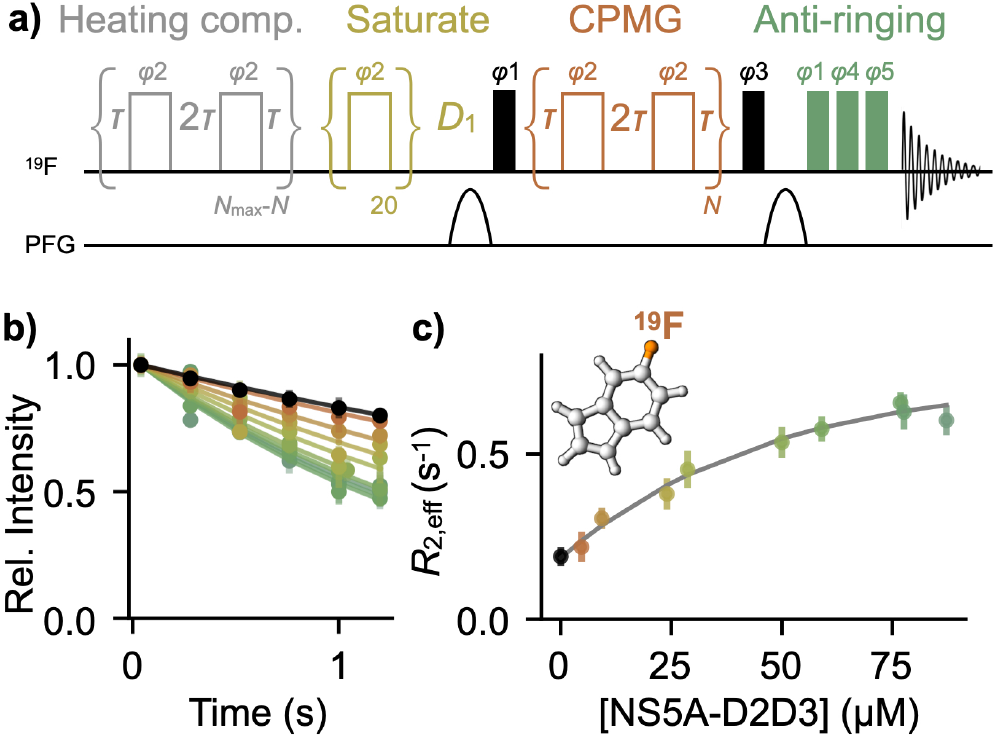
Transverse ^19^F relaxation (*R*_2,eff_) is sensitive to small-molecule binding to a disordered protein. **(a)** Modified pulse sequence for the ^19^F *R*_2,eff_ experiment.^[1]^ Narrow and wide rectangles indicate 90° and 180° pulses, respectively. The number of CPMG blocks, *N*, is varied in experiments, while the time of each block, 4*τ* is held constant. Phase cycles are reported in Supplementary Information. **(b)** Relaxation curves obtained for 50 μM 5-fluoroindole in the absence and presence of varying concentrations of NS5A-D2D3. Error bars are SEM from ≥3 technical replicates. **(c)** *R*_2,eff_ rates obtained from **b** as a function of NS5A-D2D3 concentration. Error bars represent the uncertainty in the *R*_2,eff_-fitted parameter from **b** (from the covariance matrix). A one-site binding model (grey curve), accounting for *R*_2,eff_ rates, chemical shifts, longitudinal relaxation, and translational diffusion (see text), was fit to the data, yielding an affinity constant (*K*_B_) of 260 ± 110 μM.

### Transverse spin-relaxation is the most sensitive parameter to binding

Having established that 5-fluoroindole interacts with the IDP NS5A-D2D3 allows us to assess the sensitivity of methods that are typically used within solution NMR spectroscopy to characterize small-molecule binding, such as chemical shift perturbations and changes in signal intensities, longitudinal relaxation, and translational diffusion. Firstly, protein-detected chemical shift-based experiments, such as the 2D ^1^H-^15^N HSQC (**Figures 2a, S1, S2**), are largely insensitive to the interaction between NS5A-D2D3 and 5-fluoroindole at protein:ligand ratios of 1:4 and 1:8. This holds for both protein ^1^H and ^15^N chemical shifts (**Figure 2b**,**c**) as well as changes in the intensity of protein ^1^H-^15^N cross-peaks (**Figure 2d**). It was also observed that ligand-detected NMR chemical shifts, including both ^1^H and ^19^F, are minimally sensitive to binding in a concentration-dependent manner (**Figure 2e**-**h**). Similar conclusions were drawn previously, where protein-detected NMR chemical shifts provided limited information on the binding of the small-molecule 10074-G5 to the IDP amyloid-β.^[5]^

**Figure 2.**
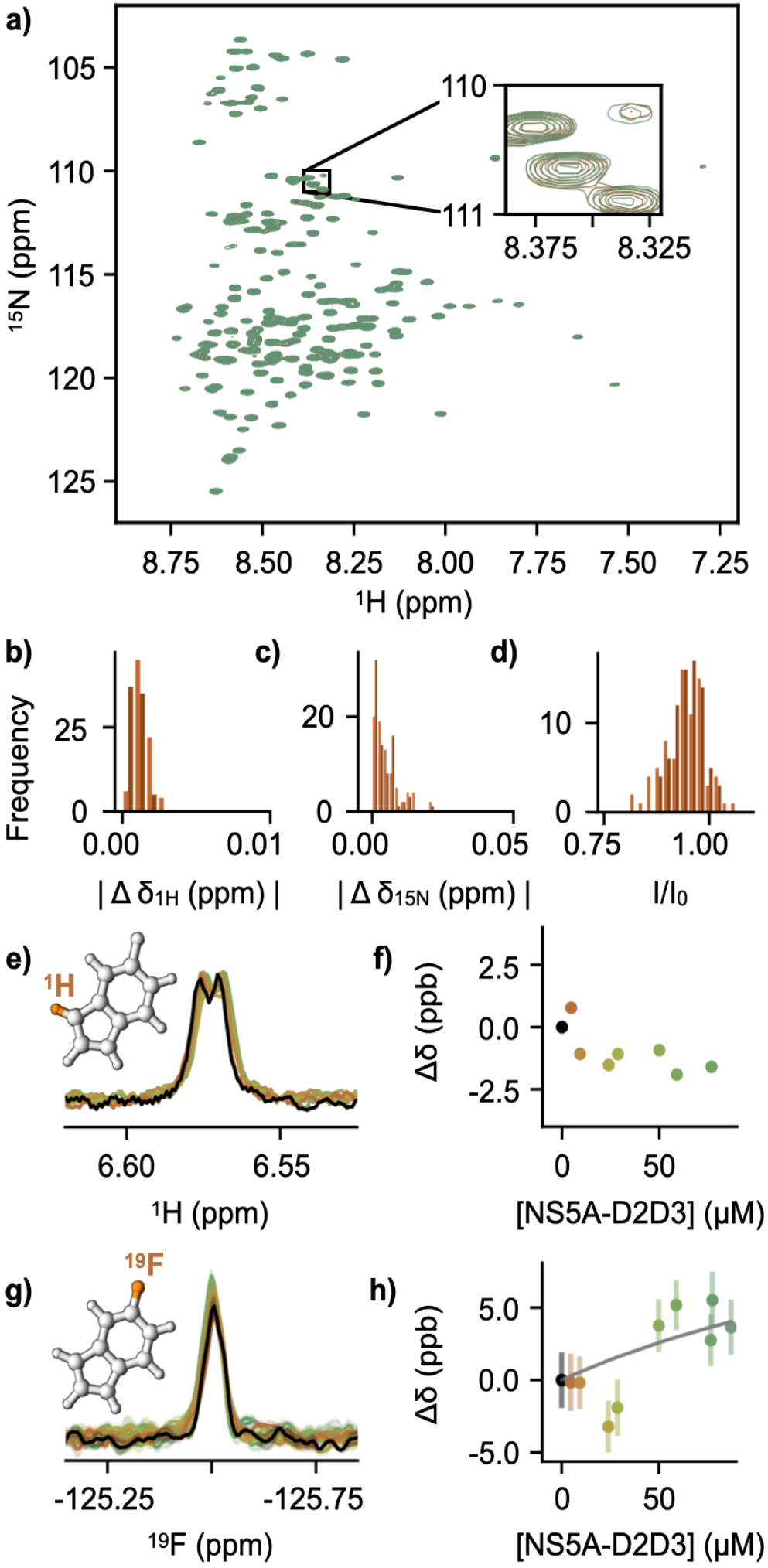
NMR chemical shifts are insensitive to IDP/small-molecule binding. **(a)** ^1^H-^15^N HSQC spectrum of 40 μM NS5A-D2D3 in the absence (green) and presence (orange) of 320 μM 5-fluoroindole acquired at 15°C. Insert shows the largest chemical shift difference calculated as shown in Supplementary Information (**Figure S1**) **(b-d)** Histogram of chemical shift changes of NS5A-D2D3 in the presence of 160 μM (light orange) and 320 μM (dark orange), relative to NS5A-D2D3 alone, showing minimal perturbations in the presence of the small molecule, including ^1^H **(b)** and ^15^N **(c)** chemical shift perturbations and intensity changes **(d)**. See also **Figure S2. (e)** Ligand-detected ^1^H chemical shifts of 50 μM 5-fluoroindole show minimal changes upon titration with NS5A-D2D3, acquired at 25°C. **(f)** Quantification of ligand-detected ^1^H chemical shift perturbations relative to 50 μM 5-fluoroindole alone, measured in parts per billion. **(g)** Ligand-detected ^19^F chemical shifts of 50 μM 5-fluoroindole show minimal changes upon titration with NS5A-D2D3, acquired at 25°C. Curves represent the average of ≥2 technical replicates. Error bars represent SEM. **(h)** Quantification of ligand-detected ^19^F chemical shift perturbations relative to 50 μM 5-fluoroindole alone, measured in parts per billion. Error bars represent SEM ≥2 technical replicates. Although the changes alone provide limited insight, together with the *R*_2,eff_ rates these shifts are important for the analysis. A one-site binding model, accounting for chemical shifts, transverse relaxation, longitudinal relaxation, and diffusion (see text), was fit the data (grey curve).

We also tested ^19^F-detected longitudinal relaxation of 50 μM 5-fluoroindole in the presence of varying concentrations of NS5A-D2D3, measured using an inversion recovery experiment, and observed minimal sensitivity to binding (**Figure S3**). Similarly, the translational diffusion of 50 μM 5-fluoroindole, measured by ^1^H Diffusion Ordered Spectroscopy (DOSY) was largely insensitive to changes in the absence and presence of 75 μM NS5A-D2D3 (**Figure S4**).

### 5-fluoroindole remains dynamic in the bound form

To gain additional insight into the interaction mechanism of 5-fluoroindole with NS5A-D2D3, a simple one-site binding model was assumed, where 5-flouroindole can exist in two states: a ‘free’ form (F) and a ‘bound’ form (B) interacting with NS5A-D2D3. The binding mechanism is likely dynamic and more complex, but here we simply assume the ‘bound’ form represents an ensemble of states all interacting with NS5A-D2D3. The increase in *R*_2,eff_ observed with increasing concentrations of NS5A-D2D3 (**Figure 1**) could arise from either an elevated intrinsic *R*_2_ of 5-fluoroindole in the bound conformation or an exchange-induced increase in *R*_2_.^[25]^ The experimental ^19^F transverse relaxation, (**Figure 1c**), ^19^F chemical shifts (**Figure 2h**), ^19^F longitudinal relaxation (**Figure S3**), and ^1^H diffusion measurements (**Figure S4**) were therefore analyzed simultaneously within the one-site binding model. In particular, we related *R*_1,F_, *R*_1,B_, *R*_2,F_, and *R*_2,B_ with free and bound rotational correlation times, *τ*_c,F_ and *τ*_c,B_, respectively via well-established equations (Supplementary Information).^[26-29]^ *τ*_c,F_, *τ*_c,B_, *k*_off_, *K*_d_, and *D*_B_, the diffusion coefficient of the bound form, were determined from a least-squares analysis; see **Experimental Procedures** (Supplementary Information). Fits obtained using all data are shown in **Figures 1c, 2h**, and **S3**. The analysis gave a *K*_d_ of 260 ± 110 μM, a *k*_off_ of 800 ± 500 s^-1^ (**Figure 3**), a *τ*_c,F_ of 27.0 ps ± 1.3 ps, a *τ*_c,B_ of 46 ps ± 10 ps, and a *D*_B_ of (1.5 ± 0.6) × 10^−9^ m^2^s^-1^. Tryptophan residues, which, like 5-fluoroindole also contain an indole motif, within disordered protein sequences have been reported to have rotational correlation components between approximately 100 and 260 ps.^[30]^ In this context, the *τ*_c,B_ we observe suggests that the small molecule remains highly dynamic in the bound state, consistent with predictions of other small molecules interacting with IDPs.^[5, 8, 11]^ Not only do ^19^F transverse relaxation rates allow for a sensitive detection of small-molecule binding to IDPs, but an analysis of the data also allows for a quantification of the associated dynamics of the interaction, dissociation constant, and off-rates. The value of the derived *K*_d_ is also of note, since the micromolar interaction observed here is the same order as often observed for lead compounds in initial drug-screening programs.^[31]^

**Figure 3.**
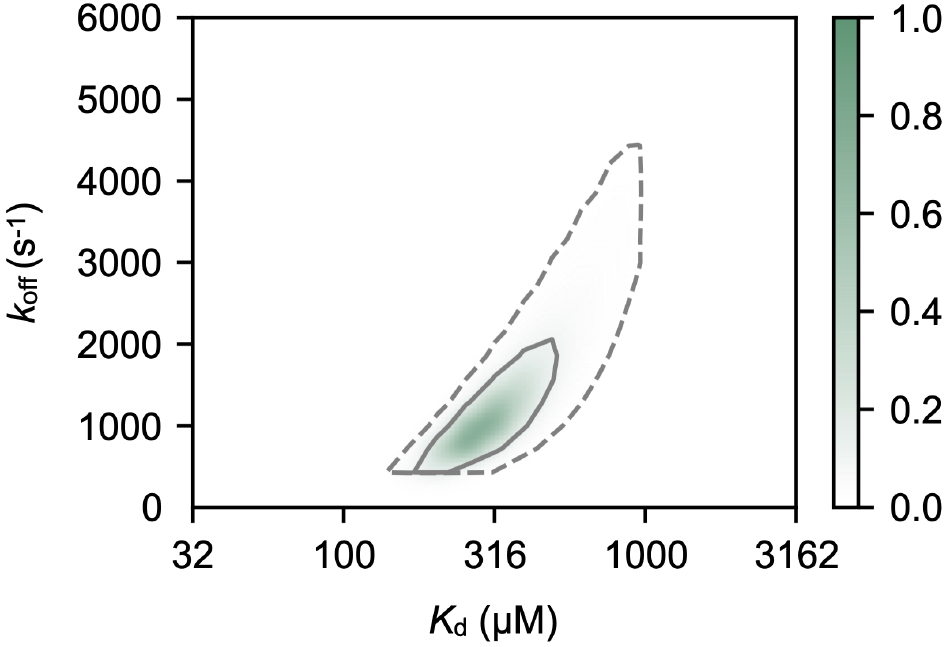
Normalized probability surface as a function of K_d_ and k_off_ show a micromolar binding affinity. The surface shown is calculated as 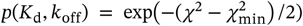, where *χ*^2^ is obtained from the least-squares fit of a one-site binding model to the experimental data. Solid and dashed lines represent 68% and 95% confidence intervals, respectively.

### When NS5A-D2D3 is less disordered, chemical shift perturbations are more significant

It has previously been reported that at high concentrations both NS5A-D2 and NS5A-D3 form transient secondary structures, potentially due to the formation of multimers.^[16]^ To confirm this also occurs for NS5A-D2D3, we performed circular dichroism (CD) measurements and observed a concentration dependent loss of disorder at and above 100 μM, suggesting transient secondary structure formation (**Figure 4a, S5**). From the CD experiments a saturation of the more structured state could not be achieved, and an equilibrium constant could therefore not be determined for the multimerization.

**Figure 4.**
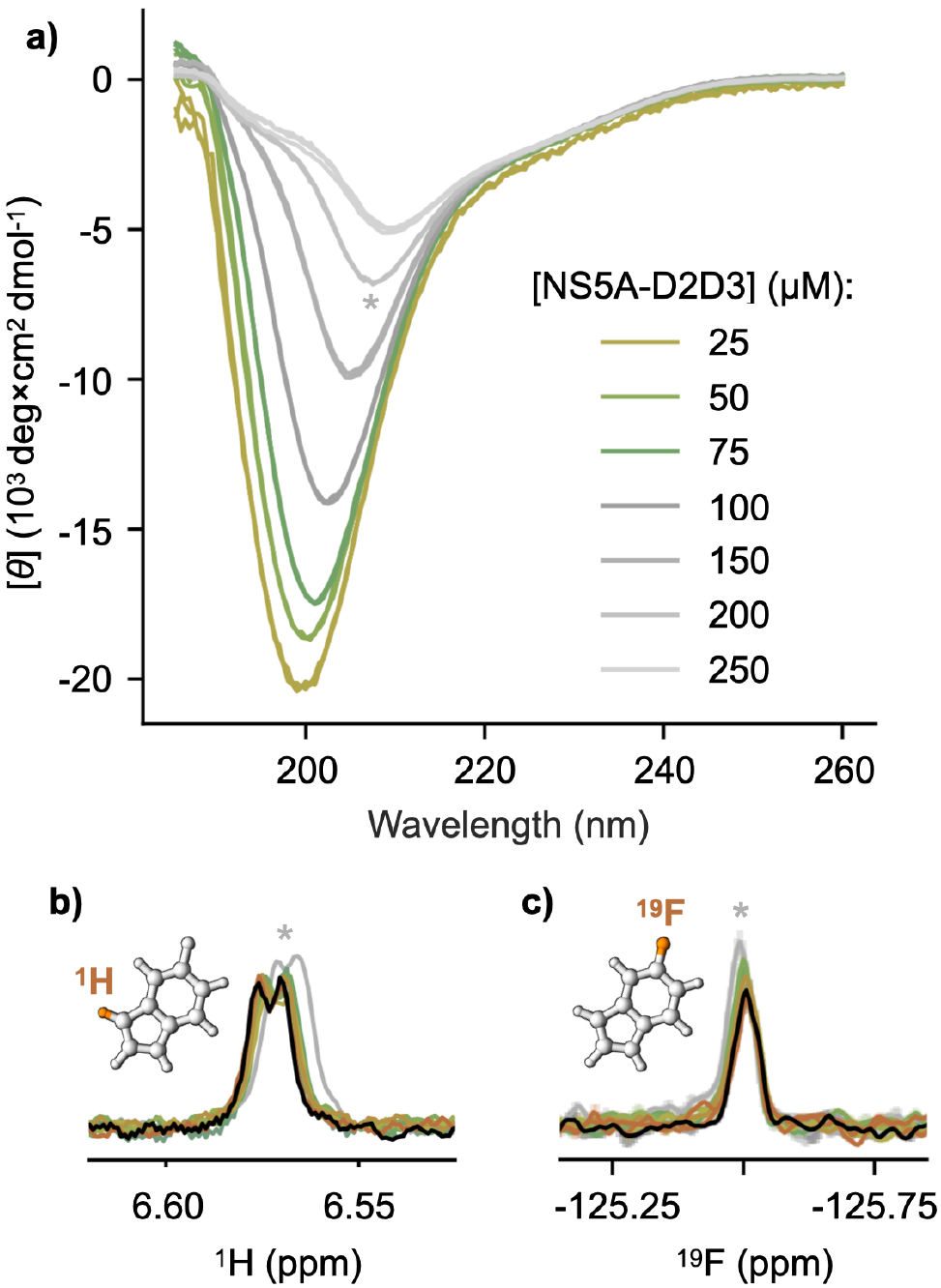
At high protein concentrations, NS5A-D2D3 becomes less disordered and shows chemical shift perturbations. **(a)** Circular dichroism measurements (molar ellipticity per residue) of various concentrations of NS5A-D2D3 demonstrating that the IDP is disordered at concentrations up until 75 μM and becomes less disordered at concentrations above 100 μM. **(b)** Ligand-detected ^1^H chemical shifts of 50 μM 5-fluoroindole from **Figure 2f** show perturbation upon titration with 200 μM NS5A-D2D3 (grey curve). Measurements were acquired at 25°C. **(c)** Ligand-detected ^19^F chemical shifts of 50 μM 5-fluoroindole from **Figure 2h** show perturbation upon titration with NS5A-D2D3 with 200 μM NS5A-D2D3 (grey curve). Measurements were acquired at 25°C.

Knowing that 5-fluoroindole interacts with NS5A-D2D3, we wondered whether this interaction alters the equilibrium of N55A-D2D3 related to a change in secondary structure propensity. At NS5A-D2D3 concentrations at or below 75 μM, we observed minimal changes in the presence of 50 μM 5-fluoroindole, consistent with the protein-detected NMR experiments (**Figure S5a, b**), that is, 5-fluoroindole generally does not alter the secondary structure propensity of NS5A-D2D3. At 200 μM NS5A-D2D3, very subtle changes were observed, suggesting that 5-fluoroindole may stabilize the less disordered state adopted by NS5A-D2D3 when it is at higher concentrations (**Figure S5c**).

Of particular interest is that the NS5A-D2D3 system at high concentrations provides a way to assess the binding of 5-fluoroindole to a less disordered state. Notably, when chemical shift perturbations of 5-fluoroindole were measured in the presence of 200 μM NS5A-D2D3, small, but significant chemical shift perturbations for both ^1^H (**Figure 4b**) and ^19^F (**Figure 4c**) were detected. This observation coincided with an increase in ^19^F longitudinal (*R*_1,eff_) and transverse (*R*_2,eff_) relaxation rates of 0.24 ± 0.02 s^-1^ and 1.40 ± 0.11 s^-1^, respectively. These data suggest a further change in the effective correlation time of 5-fluoroindole when it interacts with the less disordered state of NS5A-D2D3.

## Conclusion

NMR chemical shift perturbations are a gold-standard technique for screening and characterizing small-molecule binding to structured proteins. While significant protein-detected chemical shift perturbations have been reported for small-molecule interactions with IDPs,^[8, 31]^ these are often small, a fraction of the peak linewidths, or even undetectable in cases like the one presented here (**Figure 2**). Furthermore, it has recently been reported that HSQCs of disordered proteins are highly prone to false-positive characterization of ligand interactions due to artefacts arising from mismatched pH.^[32]^ In contrast, we report here that ligand-detected ^19^F transverse relaxation measurements are sensitive to small-molecule/IDP binding. We uncovered a micromolar binding affinity between 5-fluoroindole and NS5A-D2D3 in its disordered form, where chemical shift perturbations were minimal. We also show that the small molecule remains very dynamic when interacting with NS5A-D2D3 in its disordered state. We anticipate that ^19^F ligand-detected spin-relaxation experiments offer a promising medium-throughput screening strategy to identify small molecules that bind IDPs and other dynamic biomolecules, especially in cases where such interactions may be largely undetectable by other approaches. Moreover, the method presented is general and would also apply to measuring the transverse relaxation rate of other nuclei within ligands, such as, ^1^H, given that these can be measured accurately.

## Methods

Samples of NS5A-D2D3 were expressed and purified as described in the Supplementary Information. NMR ^19^F relaxation experiments were performed on Bruker Avance III 500 MHz (CP-prodigy probe). ^1^H-^15^N experiments were performed at 700 MHz. NMR spectra were processed and analyzed as detailed in Supplementary Information. A comprehensive list of all experiments including sample details and experimental conditions is given in the Supplementary Information.

## Supplementary information

Supplementary information^[33-43]^ includes several figures, methods, and the *R*_2,eff_ pulse program. Code that supports the findings of this study is available from GitHub at https://github.com/hansenlab-ucl/R2_IDP_small_mol. All data files are available from Zenodo at https://zenodo.org/record/7892349#.ZFKHGC8w3Uo.

## Acknowledgements

The authors acknowledge Gogulan Karunanithy and Geoff Kelly for helpful discussion. The authors also acknowledge Khushboo Matwani for assistance with protein production. GTH was supported by Schmidt Science Fellows, Rosaland Franklin Research Fellowship from Newnham College, Cambridge, and a BBSRC Discovery Fellowship (BB/X009955/1). This work was also supported by the UCL Wellcome Institutional Strategic Support Fund (204841/Z/16/Z). The BBSRC (BB/R000255/1), Wellcome Trust (ref101569/z/13/z), and the EPSRC are acknowledged for supporting the NMR facility at University College London. Access to ultra-high field NMR spectrometers was supported by the Francis Crick Institute through provision of access to the MRC Biomedical NMR Centre. The Francis Crick Institute receives its core funding from Cancer Research UK (FC001029), the UK Medical Research Council (FC001029), and the Wellcome Trust (FC001029). DFH is supported by the UKRI and EPSRC (EP/X036782/1). This paper was typeset with the bioRxiv word template by @Chrelli: www.github.com/chrelli/bioRxiv-word-template.

## Author contributions

GTH and VKS produced samples of NS5A-D2D3. GTH, AMF, and DFH collected and analyzed the data. All authors discussed the results and commented on the paper.

## References

1. J. H. Overbeck, W. Kremer, R. Sprangers, Journal of Biomolecular NMR 2020, 74, 753–766.

2. H. J. Dyson, P. E. Wright, Nature Reviews Molecular Cell Biology 2005, 6, 197–208.

3. V. Csizmok, A. V. Follis, R. W. Kriwacki, J. D. Forman-Kay, Chemical Reviews 2016, 116, 6424–6462.

4. P. Tompa, Trends in Biochemical Sciences 2002, 27, 527–533.

5. G. T. Heller, F. A. Aprile, T. C. Michaels, R. Limbocker, M. Perni, F. S. Ruggeri, B. Mannini, T. Löhr, M. Bonomi, C. Camilloni, A. De Simone, C. Felli, R. Pieratelli, T. P. J. Knowles, C. M. Dobson, M. Vendruscolo, Science Advances 2020, 6, eabb5924.

6. G. T. Heller, F. A. Aprile, M. Bonomi, C. Camilloni, A. De Simone, M. Vendruscolo, Journal of Molecular Biology 2017, 429, 2772–2779.

7. A. V. Follis, D. I. Hammoudeh, H. Wang, E. V. Prochownik, S. J. Metallo, Chemistry & Biology 2008, 15, 1149–1155.

8. P. Robustelli, A. Ibanez-de-Opakua, C. Campbell-Bezat, F. Giordanetto, S. Becker, M. Zweckstetter, A. C. Pan, D. E. Shaw, Journal of the American Chemical Society 2022, 144, 2501–2510.

9. G. T. Heller, P. Sormanni, M. Vendruscolo, Trends in Biochemical Sciences 2015, 40, 491–496.

10. F. Jin, C. Yu, L. Lai, Z. Liu, PLoS Computational Biology 2013, 9, e1003249.

11. T. Löhr, K. Kohlhoff, G. T. Heller, C. Camilloni, M. Vendruscolo, ACS Chemical Neuroscience 2022, 13, 1738–1745.

12. F. E. Thomasen, K. Lindorff-Larsen, Biochemical Society Transactions 2022, 50, 541–554.

13. S. Bottaro, K. Lindorff-Larsen, Science 2018, 361, 355–360.

14. M. Bonomi, G. T. Heller, C. Camilloni, M. Vendruscolo, Current Opinion in Structural Biology 2017, 42, 106–116.

15. S. Rauscher, V. Gapsys, M. J. Gajda, M. Zweckstetter, B. L. De Groot, H. Grubmüller, Journal of Chemical Theory and Computation 2015, 11, 5513–5524.

16. A. Badillo, V. Receveur-Brechot, S. Sarrazin, F.-X. Cantrelle, F. Delolme, M.-L. Fogeron, J. Molle, R. Montserret, A. Bockmann, R. Bartenschlager, V. Lohmann, G. Lippens, S. Ricard-Blum, X. Hanoulle, F. Penin, Biochemistry 2017, 56, 3029–3048.

17. M. Dujardin, V. Madan, R. Montserret, P. Ahuja, I. Huvent, H. Launay, A. Leroy, R. Bartenschlager, F. Penin, G. Lippens, Journal of Biological Chemistry 2015, 290, 19104–19120.

18. X. Hanoulle, A. Badillo, J.-M. Wieruszeski, D. Verdegem, I. Landrieu, R. Bartenschlager, F. Penin, G. Lippens, Journal of Biological Chemistry 2009, 284, 13589–13601.

19. G. Heller, L. Yu, D. Hansen, in NMR Spectroscopy for Probing Functional Dynamics at Biological Interfaces, Royal Society of Chemistry, 2022, pp. 383–410.

20. I. C. Felli, R. Pierattelli, Intrinsically disordered proteins studied by NMR spectroscopy, Vol. 870, Springer, 2015.

21. M. P. Williamson, Progress in Nuclear Magnetic Resonance Spectroscopy 2013, 73, 1–16.

22. D. Ban, L. I. Iconaru, A. Ramanathan, J. Zuo, R. W. Kriwacki, Journal of the American Chemical Society 2017, 139, 13692–13700.

23. S. Meiboom, D. Gill, Review of Scientific Instruments 1958, 29, 688–691.

24. H. Y. Carr, E. M. Purcell, Physical Review 1954, 94, 630.

25. H. M. McConnell, The Journal of Chemical Physics 1958, 28, 430–431.

26. C. D. Kroenke, J. P. Loria, L. K. Lee, M. Rance, A. G. Palmer, Journal of the American Chemical Society 1998, 120, 7905–7915.

27. A. Abragam, The Principles of Nuclear Magnetism, Oxford university press, 1961.

28. M. Lu, R. Ishima, T. Polenova, A. M. Gronenborn, Journal of Biomolecular NMR 2019, 73, 401–409.

29. M. Lu, S. Sarkar, M. Wang, J. Kraus, M. Fritz, C. M. Quinn, S. Bai, S. T. Holmes, C. Dybowski, G. P. Yap, J. Struppe, I. V. Sergeyev, W. Maas, A. M. Gronenborn, T Polenova, The Journal of Physical Chemistry B 2018, 122, 6148–6155.

30. N. Jain, D. Narang, K. Bhasne, V. Dalal, S. Arya, M. Bhattacharya, S. Mukhopadhyay, Biophysical Journal 2016, 111, 768–774.

31. L. I. Iconaru, D. Ban, K. Bharatham, A. Ramanathan, W. Zhang, A. A. Shelat, J. Zuo, R. W. Kriwacki, Scientific Reports 2015, 5, 1–16.

32. A. K. Pandey, C. R. Buchholz, N. Nathan Kochen, W. C. Pomerantz, A. R. Braun, J. N. Sachs, ACS Chemical Neuroscience 2023, 14, 800–808.

33. R. L. Vold, The Journal of Chemical Physics 1972, 56, 3210–3216.

34. E. O. Stejskal, J. E. Tanner, The Journal of Chemical Physics 1965, 42, 288–292.

35. A. Chen, D. Wu, C. S. Johnson Jr, Journal of the American Chemical Society 1995, 117, 7965–7970.

36. F. Delaglio, S. Grzesiek, G. W. Vuister, G. Zhu, J. Pfeifer, A. Bax, Journal of Biomolecular NMR 1995, 6, 277–293.

37. J. J. Helmus, C. P. Jaroniec, Journal of Biomolecular NMR 2013, 55, 355–367.

38. M. Newville, T. Stensitzki, D. B. Allen, M. Rawlik, A. Ingargiola, A. Nelson, Astrophysics Source Code Library 2016, ascl: 1606.1014.

39. W. Lee, M. Tonelli, J. L. Markley, Bioinformatics 2015, 31, 1325–1327.

40. N. Tjandra, A. Szabo, A. Bax, Journal of the American Chemical Society 1996, 118, 6986–6991.

41. G. Pagès, S. V. Dvinskikh, I. Furó, Journal of Magnetic Resonance 2013, 234, 35–43.

42. D. F. Hansen, P. Vallurupalli, P. Lundström, P. Neudecker, L. E. Kay, Journal of the American Chemical Society 2008, 130, 2667–2675.

43. D. Verdegem, A. Badillo, J.-M. Wieruszeski, I. Landrieu, A. Leroy, R. Bartenschlager, F. Penin, G. Lippens, X. Hanoulle, Journal of Biological Chemistry 2011, 286, 20441–20454.

