## Supplementary Information for "Small-molecule binding to an intrinsically disordered protein revealed by experimental NMR ^19^F transverse spin-relaxation"

bioRxiv

### Experimental Procedures

#### Sample preparation

A recombinant construct of the D2 and D3 domains of HCV NS5A (NS5A-D2D3, residues 247-466) was purified as follows:[1-3] the codon-optimized synthetic sequence coding for domains 2 and 3 of the HCV NS5A protein from the JFH1 strain (GenBank™ accession number AB047639, genotype 2a, purchased from Genscript) was introduced into the bacterial expression vector pET-28a(+) with a preceding 6x-His tag followed by a Tobacco Etch Virus (TEV) cleavage site at the N-terminus. *Escherichia coli* (*E. coli*) OverExpress C41(DE3) cells were transformed with this vector and grown at 37°C in M9 media with isotopically enriched $^{15}$N $NH_4Cl$ (1 g/L) as the sole nitrogen source or lysogeny broth (LB) media to obtain labelled and unlabeled protein, respectively. Expression was induced at an $OD_{600}$ between 0.6 and 0.8 by addition of 0.5 mM isopropyl β-D-thiogalactopyranoside (IPTG) and the cells were left shaking for 3-4 hours at 37°C. Cells were harvested by centrifugation. Pellets from 1L culture growths were dissolved in 50 mL of 100 mM $NaH_2PO_4$, 10 mM Tris-HCl, 20 mM imidazole, 200 mM NaCl, 2mM β-mercaptoethanol, pH 8.0 with the addition of Roche protease inhibitors tablets. Dissolved pellets were boiled at 120°C until sedimentation was observed. Pellets were vortexed and flash frozen in $LN_2$. After melting, small amounts of DNAse were added before centrifugation for 1 h at 27k RCF at 5°C. The supernatant was filtered through 0.8 um filter before being loaded onto a Ni-NTA column, washed with matching buffer containing 2M NaCl to remove nucleic acids, and eluted with a linear imidazole gradient (20–250 mM) in the absence of high salt. The protein was then concentrated and injected into a Superdex 75 column (GE Healthcare) and subjected to size-exclusion chromatography at 5°C. The His tag of pure fractions was cleaved by a TEV protease (containing its own 6xHis tag) at room temperature for 2 h. His beads were added to remove the TEV protease and uncleaved protein before a final size- exclusion step while exchanging the buffer into 25 mM Tris, 150 mM NaCl, 1 mM tris(2-carboxyethyl)phosphine (TCEP), pH 7.0. Protein was concentrated using Amicon Ultra Centrifugal filters with a 3KDa cutoff.

5-fluoroindole was purchased from Sigma-Aldrich (CAS Number: 399-52-0), dissolved in DMSO-d6 at 1 M concentration and kept frozen at -20°C until use.

All NMR samples were prepared in 25 mM Tris, 150 mM NaCl, 1 mM TCEP, pH 7.0, 2% $D_2O$.

#### NMR Spectroscopy

$^{1}$H-ligand detected chemical shifts and all $^{19}$F NMR measurements were performed on a 11.7 T Bruker AVANCE III spectrometer, equipped with a Prodigy H&F-C/N-D TCI cryoprobe. The temperature for all ligand-detected measurements was 298K. $^{1}$H and $^{19}$F chemical shifts were referenced with respect to 4,4-dimethyl-4-silapentane-1-sulfonic acid (DSS) and trichloro-fluoro-methane ($CFCl_3$), respectively. 1D $^{1}$H spectra were acquired with the standard zgesgp Bruker pulse sequence with 8,192 complex points, a spectral width of 7,500 Hz, and an acquisition time of 1 s. 640 scans were collected per experiment. 1D $^{19}$F spectra were acquired using the aring pulse sequence to minimize acoustic ringing. Spectra were obtained with 5,120 complex points, a spectral width of 46,875 Hz, and an acquisition time of 0.1 s. 3,200 scans were collected per experiment. The $^{19}$F frequency carrier was centered at −120 ppm. Effective $^{19}$F longitudinal (spin-lattice) relaxation rates, $R_{1,\mathrm{eff}}$, were measured using an inversion-recovery[4] experiment with a recycling delay of 16 s and relaxation delays of 0.078125, 0.15625, 0.3125, 0.625, 1.25, 2.5, 5, 10, 20 s. These experiments were acquired with 16,384 complex points, an acquisition time of 0.2 s, 32 scans per increment, and a spectral width of 75,000 Hz. The $^{19}$F frequency carrier was centered at −125 ppm. Measurements were repeated at least in duplicate to estimate the experimental error. Effective $^{19}$F transverse (spin-spin) relaxation rates, $R_{2,\mathrm{eff}}$, were measured using a CPMG-based $R_2$ experiment (**Figure 1a,** see also **$R_{2,\mathrm{eff}}$ Pulse Program**) with a relaxation delay ($D_1$) of 2 s. The number, $N$, of CPMG blocks ($\tau_{\mathrm{CPMG}} - \pi - 2\tau_{\mathrm{CPMG}} - \pi - \tau_{\mathrm{CPMG}}$) was varied between experiments, such that $N =$ 1, 7, 13, 19, 25, or 30, where $\tau_{\mathrm{CPMG}}$ was held constant at 10 ms across all experiments. A heating compensation block was also included before the start of the experiment (**Figure 1a,** see also **$R_{2,\mathrm{eff}}$ Pulse Program).** $R_{2,\mathrm{eff}}$ experiments were acquired with 1,024 complex points, an acquisition time of 0.11 s, 64 scans per increment, and a spectral width of 9,398 Hz. The $^{19}$F frequency carrier was centered at −120 ppm. Measurements were repeated at least in triplicate to estimate the experimental error.

2D $^{1}$H-$^{15}$N HSQC spectra of NS5A-D2D3 were collected on a 16.5 T Bruker Avance III 700 MHz spectrometer, equipped with a TCI cryoprobe. The experiments were acquired with 1,024 and 128 complex points in the $^{1}$H and $^{15}$N dimensions with spectral widths of 11,161 and 2,484 Hz, respectively. The frequency carrier was centered at 4.693 and 121.500 ppm in the $^{1}$H and $^{15}$N dimensions, with acquisition times of 0.09 and 0.05 s, respectively. 64 scans were collected per experiment. The temperature for all protein-detected 2D measurements was 288K.

$^{1}$H Diffusion Ordered SpectroscopY (DOSY) measurements[5] of 5-fluoroindole were measured on a 18.8 T Bruker Avance III 800 MHz spectrometer equipped with a TCI cryoprobe at 298K. Measurements were performed with the standard ledbpgppr2s Bruker pulse sequence[6] using bipolar gradients and solvent suppression. DOSY experiments were acquired with 16,384 complex points, an acquisition time of 1.3 s, 16 scans per increment, and a spectral width of 12,821 Hz. We used a diffusion time ($\Delta$) of 200 ms, a gradient pulse length ($\delta$) of 2 ms with a smoothed rectangular shape, and a time, $\tau$, to phase/rephase bipolar gradients of 0.2 ms. We collected 16 gradient strengths, $g$, linearly spaced between 0.681 and 20.434 G cm$^{-1}$.

All data were processed and analyzed using nmrPipe[7], nmrGlue[8], and lmfit[9]. 2D data were visualized using Sparky.[10] 1D data were fit to Lorentzian curves and relaxation rates were obtained by fitting peak intensities to single exponential functions using in-house software written in Python, available on GitHub (https://github.com/hansenlab-ucl/R2_IDP_small_mol).

DOSY spectra were fit to the following equation:

$$\frac{I}{I_0} = e^{-D\,\gamma_{\rm H}^2 g^2 \delta^2\left(\Delta - \frac{\delta}{3} - \frac{\tau}{2}\right)},$$

where $I$ is the observed intensity, $I_0$ is the intensity of the unattenuated signal, $D$ is the diffusion coefficient, and $\gamma_{\rm H}$ is the gyromagnetic ratio of $^{1}$H.

### Analysis of relaxation rates, translational diffusion, and chemical shifts

We used a least-squares fitting analysis to assess the longitudinal and transverse relaxation rates, chemical shifts, and translational diffusion. We assumed a simple two-site exchange model in fast-to-intermediate exchange between a free (F) and bound (B) state of 5-fluoroindole, that is,

$$\mathrm{F} \underset{k_{\rm bf}}{\overset{k_{\rm fb}}{\rightleftharpoons}} \mathrm{B} \qquad (1)$$

in which $k_{\rm fb}$ and $k_{\rm bf}$ are the pseudo-first order forward and reverse reaction rate constants, respectively, such that $k_{\rm fb} = k_{\rm on} * [P]$ and $k_{\rm bf} = k_{\rm off}$, where $k_{\rm on}$ and $k_{\rm off}$ are the second-order binding association rate and the dissociation, respectively.

The intrinsic relaxation rates $R_{1,\rm F}$, $R_{1,\rm B}$, $R_{2,\rm F}$, and $R_{2,\rm B}$, were described in terms of rotational correlation times, $\tau_{\rm c}$, using the $^{1}$H-$^{19}$F two-spin system approximation[11] accounting for the dipole-dipole interaction of fluorine with the two nearest hydrogens and the $^{19}$F chemical shift anisotropy (CSA) contribution according to the equations below:

$$R_1 = \frac{d^2}{2}\left[3J(\omega_{\rm F}) + J(\omega_{\rm H} - \omega_{\rm F}) + 6J(\omega_{\rm H} + \omega_{\rm F}) + c^2 J(\omega_{\rm F})\right]$$

$$R_2 = \frac{d^2}{4}\left[4J(0) + 3J(\omega_{\rm F}) + J(\omega_{\rm H} - \omega_{\rm F}) + 6J(\omega_{\rm H}) + 6J(\omega_{\rm H} + \omega_{\rm F})\right] + \frac{c^2}{6}\left(4J(0) + 3J(\omega_{\rm F})\right)$$

where the power spectral density function, $J(\omega)$, describing the frequency distribution of stochastic motion that modulate the dipole-dipole and CSA Hamiltonians[12-13] such that $J^{\rm Dipolar}(\omega) \approx J^{\rm CSA}(\omega) = J(\omega)$ is given by

$$J(\omega) = \frac{2}{5}\frac{\tau_c}{1 + \tau_c^2\omega^2}$$

and $\omega$ is the angular frequency. Furthermore, $d = \frac{\mu_0 h \gamma_{\rm H}\gamma_{\rm F}}{8\pi^2 2 r_{\rm FH}^3}$ and $c = \frac{\gamma_{\rm F} B_0 \Delta\sigma}{\sqrt{3}}$, $\mu_0$ is the permeability of free space, $h$ is Planck's constant, $\gamma_{\rm H}$ and $\gamma_{\rm F}$ are the gyromagnetic ratios for $^{1}$H and $^{19}$F, respectively, and $B_0$ is the static magnetic field strength. $r_{\rm FH}$, the distance between the two nuclei for both $^{19}$F-$^{1}$H pairs, was approximated as 2.6 Å.[14] $\Delta\sigma = \sigma_{||} - \sigma_{\perp}$ where $\sigma_{||}$ and $\sigma_{\perp}$ are the principle components of the $^{19}$F CSA tensor. $\Delta\sigma$ was approximated as 52.1 ppm.[14-15] The reduced anisotropy, $\eta$, was taken to be 0.5. [14-15]

The NMR relaxation rates of free and bound 5-fluoroindole above were used as input for evolution matrices to account for the effects of chemical exchange.[16-17] The evolution of the transverse $^{19}$F magnetization, used to model both the transverse relaxation rate, $R_{2,\rm eff}$, and the chemical shifts was described by the evolution matrix, $\mathbf{\Gamma}_{\perp}$:

$$\mathbf{\Gamma}_{\perp} = \begin{bmatrix} -R_{2,\rm F} - k_{\rm fb} & k_{\rm bf} \\ k_{\rm fb} & -R_{2,\rm B} + \mathrm{i}\Delta\omega - k_{\rm bf} \end{bmatrix}, \qquad (2)$$

where $R_{2,\mathrm{F}}$ and $R_{2,\mathrm{B}}$ are the intrinsic transverse relaxation rates of the free and bound species, respectively, and $\Delta\omega$ is the difference in chemical shift between the two species. The time-evolution of the transverse magnetization ($\mathrm{I}_{\mathrm{xy},j} = \mathrm{I}_{\mathrm{x},j} + i\mathrm{I}_{\mathrm{y},j}$ where $j \in \{\mathrm{F}, \mathrm{B}\}$ and $i$ is the imaginary unit) is described by the following homogeneous differential equation:

$$\frac{d}{dt}\begin{bmatrix}\mathrm{I}_{\mathrm{xy,F}}\\ \mathrm{I}_{\mathrm{xy,B}}\end{bmatrix} = \mathbf{\Gamma}_{\perp}\begin{bmatrix}\mathrm{I}_{\mathrm{xy,F}}\\ \mathrm{I}_{\mathrm{xy,B}}\end{bmatrix} \quad (3)$$

$$\frac{d}{dt}\mathbf{I}_{\mathrm{xy}} = \mathbf{\Gamma}_{\perp}\mathbf{I}_{\mathrm{xy}} \quad (4)$$

and the time-evolution therefore given by

$$\mathbf{I}_{\mathrm{xy}}(\mathrm{t}) = \mathbf{I}_{\mathrm{xy},0}\exp(\,\mathbf{\Gamma}_{\perp}t) \quad (5)$$

For free-precession experiments, the observed transverse relaxation rate, $R_{2,\mathrm{eff}}$, is the smallest of the real part of the eigenvalues of $\mathbf{\Gamma}_{\perp}$, while the observed chemical shift, $\delta_{\mathrm{eff}}$, is the imaginary part of this eigenvalue. Transverse relaxation rates observed in CPMG experiments were modelled by successive free-precession of $\tau_{\mathrm{CPMG}}$ and 180º pulses, as described previously.[18]

The evolution of the longitudinal $^{19}$F magnetization ($\mathrm{I}_{\mathrm{z,j}}$, where $j \in \{\mathrm{F}, \mathrm{B}\}$) used to model the longitudinal relaxation rate, $R_{1,\mathrm{eff}}$, from the inversion recovery (IR) experiments was described by the evolution matrix, $\mathbf{\Gamma}_{\parallel,\mathrm{IR}}$.

$$\frac{d}{dt}\begin{bmatrix}1\\ \mathrm{I}_{\mathrm{z,F}}\\ \mathrm{I}_{\mathrm{z,B}}\end{bmatrix} = \mathbf{\Gamma}_{\parallel,\mathrm{IR}}\begin{bmatrix}1\\ \mathrm{I}_{\mathrm{z,F}}\\ \mathrm{I}_{\mathrm{z,B}}\end{bmatrix}, \text{ where}$$

$$\mathbf{\Gamma}_{\parallel,\mathrm{IR}} = \begin{bmatrix} 0 & 0 & 0 \\ R_{1,\mathrm{F}}\left(\frac{k_{\mathrm{bf}}}{k_{\mathrm{fb}} + k_{\mathrm{bf}}}\right) & -k_{\mathrm{fb}} - R_{1,\mathrm{F}} & k_{\mathrm{bf}} \\ R_{1,\mathrm{B}}\left(\frac{k_{\mathrm{fb}}}{k_{\mathrm{fb}} + k_{\mathrm{bf}}}\right) & k_{\mathrm{fb}} & -k_{\mathrm{b}} - R_{1,\mathrm{B}} \end{bmatrix}$$

The effective translational diffusion (TD) rates observed under chemical exchange [17], were described by

$$\frac{d}{dt}\begin{bmatrix}\mathrm{I}_{\mathrm{z,F}}\\ \mathrm{I}_{\mathrm{z,B}}\end{bmatrix} = \mathbf{\Gamma}_{\parallel,\mathrm{TD}}\begin{bmatrix}\mathrm{I}_{\mathrm{z,F}}\\ \mathrm{I}_{\mathrm{z,B}}\end{bmatrix}, \text{ where}$$

$$\mathbf{\Gamma}_{\parallel,\mathrm{TD}} = \begin{bmatrix} -(2\pi q)^2 D_{\mathrm{F}} - k_{\mathrm{fb}} - R_{1,\mathrm{F}} & k_{\mathrm{bf}} \\ k_{\mathrm{fb}} & -(2\pi q)^2 D_{\mathrm{B}} - k_{\mathrm{b}} - R_{1,\mathrm{B}} \end{bmatrix} \text{ and}$$

where $R_{1,\mathrm{F}}$ and $R_{1,\mathrm{B}}$ are the longitudinal relaxation rates of the free and bound species and

$$q = \frac{\gamma g \delta}{2\pi}.$$

For the least-squares fit the following cost-function was minimized

$$\chi^2(k_{\mathrm{off}}, \mathrm{K_d}, \Delta\omega, \tau_{\mathrm{c,B}}) = \frac{\left(R_{2,\mathrm{eff}}^{\mathrm{calc}} - R_{2,\mathrm{eff}}^{\mathrm{obs}}\right)^2}{\sigma\left(R_{2,\mathrm{eff}}^{\mathrm{obs}}\right)} + \frac{\left(\delta_{\mathrm{eff}}^{\mathrm{calc}} - \delta_{\mathrm{eff}}^{\mathrm{obs}}\right)^2}{\sigma\left(\delta_{\mathrm{eff}}^{\mathrm{obs}}\right)} + \frac{\left(D_{\mathrm{eff}}^{\mathrm{calc}} - D_{\mathrm{eff}}^{\mathrm{obs}}\right)^2}{\sigma\left(D_{\mathrm{eff}}^{\mathrm{obs}}\right)} + \frac{\left(R_{1,\mathrm{eff}}^{\mathrm{calc}} - R_{1,\mathrm{eff}}^{\mathrm{obs}}\right)^2}{\sigma\left(R_{1,\mathrm{eff}}^{\mathrm{obs}}\right)}$$

Final parameters were $k_{\mathrm{off}} = 800 \pm 500\ \mathrm{s}^{-1}$, $K_{\mathrm{d}} = 260 \pm 110\ \mu\mathrm{M}$, $\Delta\omega$=0.016 ± 0.006 ppm, $\tau_{\mathrm{c,F}} = 27.0 \pm 1.3$ ps, $\tau_{\mathrm{c,B}} = 46 \pm 10$ ps, and $D_{\mathrm{B}}$= 1 (1.5 ± 0.6) × 10^-9 $\mathrm{m}^2\mathrm{s}^{-1}$.

### Circular dichroism

CD spectra of NS5A-D2D3 at various concentrations were recorded on a Chirascan dichrograph (AppliedPhotophysics). Measurements were carried out at room temperature in a 0.1 mm path length quartz cuvette. Spectra were recorded in triplicate between 185 and 260 nm with a 0.5 nm increment and a 2 s integration time. Spectra were processed and baseline corrected using the Chirascan software. mDeg were converted to units of molar ellipticity per residue. All measurements were performed in 25 mM Tris, 150 mM NaCl, 1 mM TCEP, pH 7.0.

a)

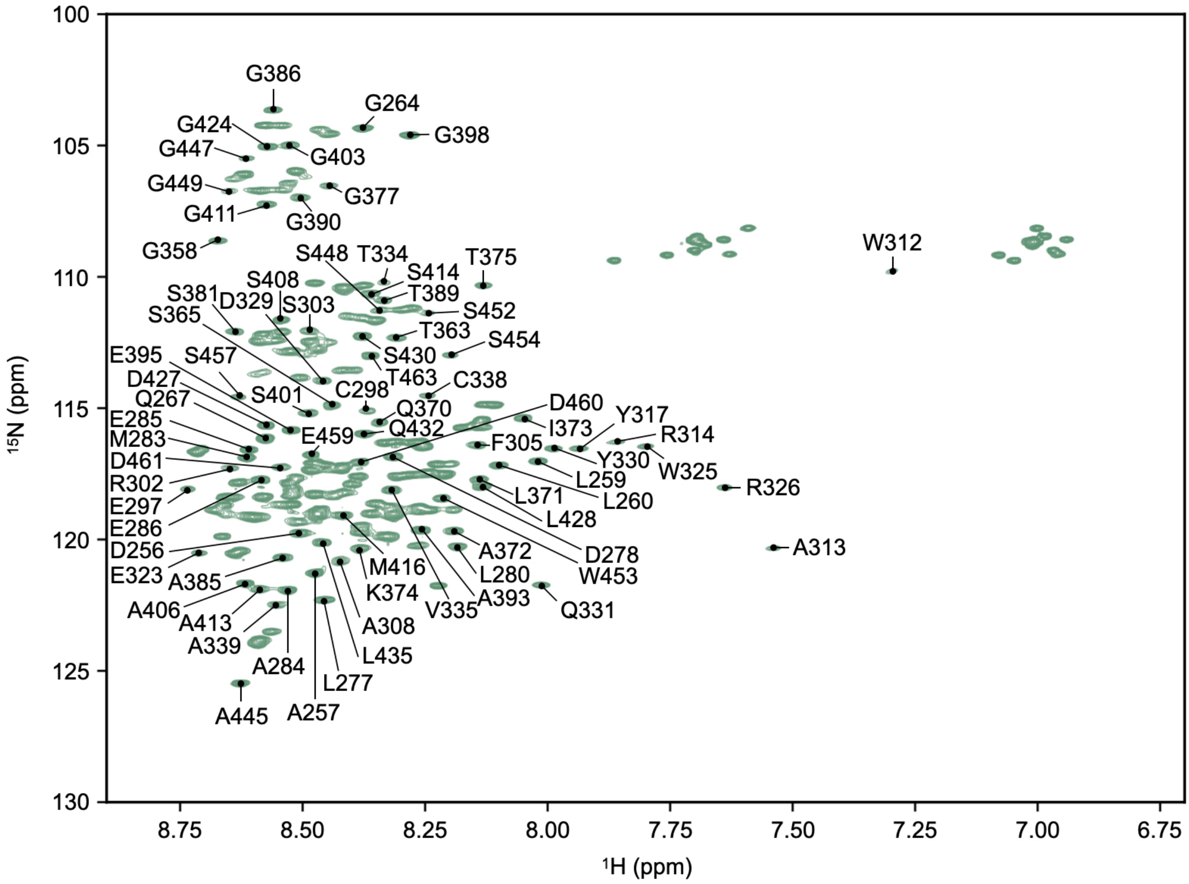


**Figure S1. Chemical shift assignments for NS5A-D2D3.** $^{1}$H-$^{15}$N HSQC spectrum of 40 µM NS5A-D2D3 acquired at 15°C with chemical shift assignments shown for the residues used in the analysis (**Figures 2**, **S2**). Assignments were taken from BMRB entries 16165[2] and 16798.[19] Overlapping and ambiguous peaks (not labelled) were discarded from the analysis.

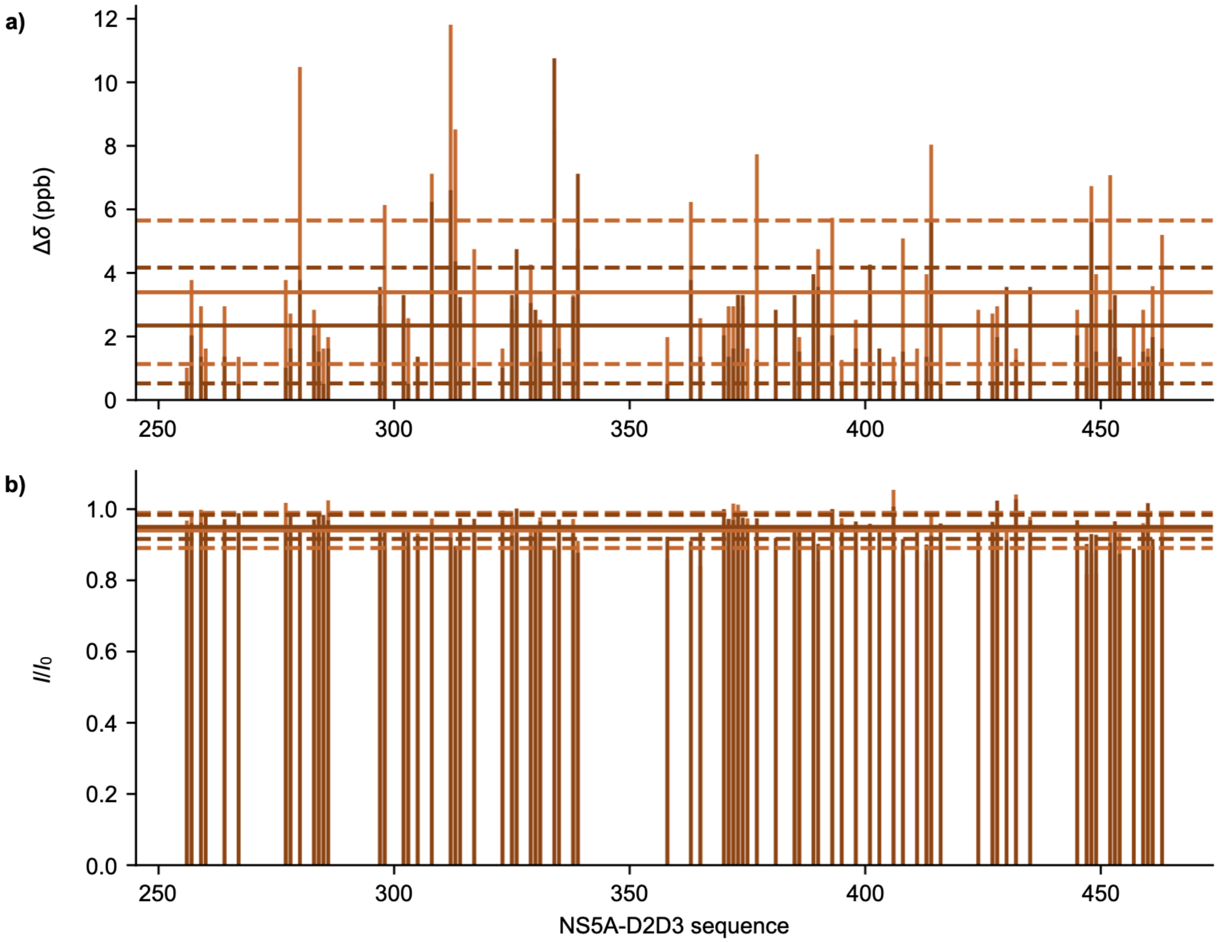


**Figure S2. Protein-detected changes upon binding 5-fluoroindole.** Sequence-dependent changes in the $^1$H-$^{15}$N HSQC spectra of NS5A-D2D3 in the presence of 160 µM (light orange) and 320 µM (dark orange) 5-fluoroindole, relative to NS5A-D2D3 alone. **(a)** Changes in chemical shifts, calculated according to $\Delta\delta = \sqrt{\frac{1}{B_{1\mathrm{HN}}}(\Delta\delta_{1\mathrm{H}})^2 + \frac{1}{B_{15\mathrm{N}}}(\Delta\delta_{15\mathrm{N}})^2}$ where $\Delta\delta_{1\mathrm{H}}$ and $\Delta\delta_{15\mathrm{N}}$ are the respective $^1$H and $^{15}$N chemical shift differences between the apo and holo states and $B_{1\mathrm{HN}} = 0.628$ and $B_{15\mathrm{N}} = 3.865$ are scaling factors, such that $B_{i\in\{1\mathrm{HN},15\mathrm{N}\}} = \sqrt{\frac{\sum \sigma^2_{\mathrm{AA}}}{n}}$, where $\sigma^2_{\mathrm{AA}}$ is the SD of a nucleus type ($^1$H$^N$ or $^{15}$N) for a standard, non-proline residue, and $n$ is the number of data points as reported in the database of chemical shifts (http://www.bmrb.wisc.edu). **(b)** Sequence-dependent changes in intensity, relative to NS5A-D2D3 alone. Intensities of each spectrum were normalized by the same peak (A393) to account for small differences. In both panels, solid lines correspond to average values across the sequence while dashed lines are ± one SD. Measurements were acquired at 15°C.

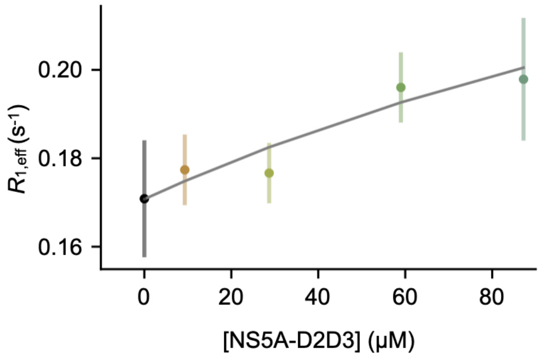


**Figure S3. $R_1$ relaxation rates of 5-fluoroindole in the presence of NS5A-D2D3.** $R_1$ relaxation rates of 5-fluoroindole measured using the $^{19}F$ inversion recovery experiment[4] in the absence and presence of various concentrations of NS5A-D2D3. A one-site binding model, accounting for chemical shifts, transverse relaxation, longitudinal relaxation, and diffusion (see **Analysis of relaxation rates, translational diffusion, and chemical shifts**), was fit the data (grey curve). Error bars represent the uncertainty in the $R_{1,eff}$ −fitted parameter (from the covariance matrix of the exponential fit to intensities from the inversion recovery experiment).

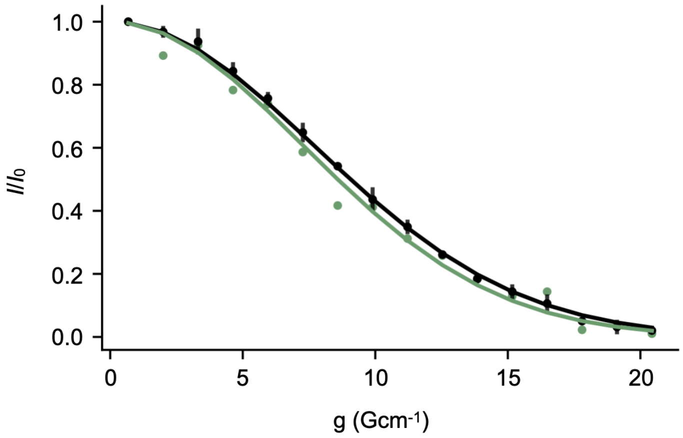


**Figure S4. Ligand-detected $^1H$ Diffusion Ordered SpectroscopY shows minimal changes in the presence of NS5A-D2D3.** $^1H$ DOSY decays shown for 50 µM 5-fluoroindole in the presence (green points) and absence (black points) of 75 µM NS5A-D2D3. Solid lines show fits to the data. Data were collected at 25°C. Error bars on the apo sample represent SD of three technical replicates.

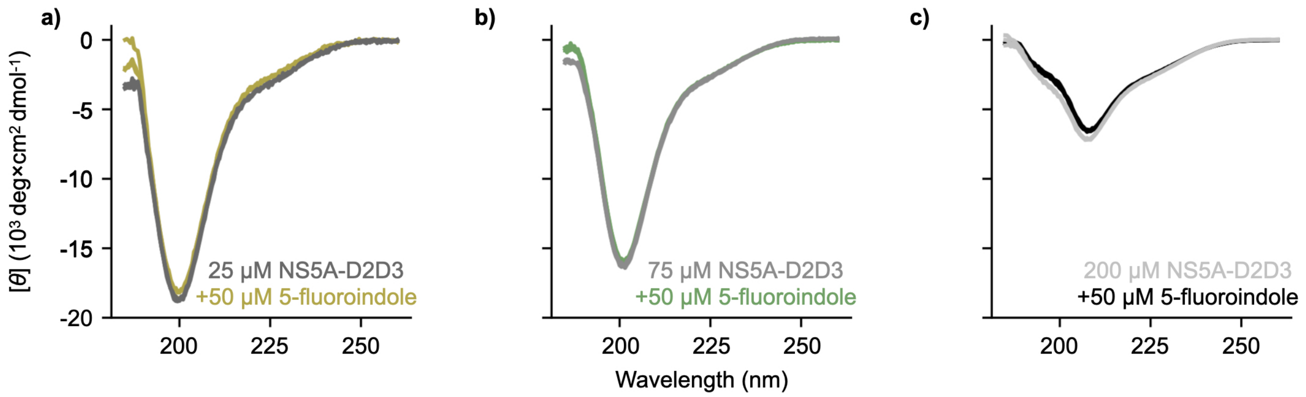


**Figure S5. Circular dichroism measurements of NS5A-D2D3 and 5-fluoroindole.** Circular dichroism measurements (molar ellipticity per residue) of NS5A-D2D3 at 25 **(a)**, 75 **(b)**, and 200 **(c)** µM in the presence (color) and absence (grey) of 50 µM 5-fluoroindole. Minimal changes in the secondary structural content are observed at 25 and 75 µM protein concentration. At 200 µM NS5A-D2D3, subtle changes in the protein secondary structural content are observed. Measurements were performed at 25°C. Three technical replicates are shown.

### References

1. A. Badillo, V. Receveur-Brechot, S. Sarrazin, F.-X. Cantrelle, F. Delolme, M.-L. Fogeron, J. Molle, R. Montserret, A. Bockmann, R. Bartenschlager, V. Lohmann, G. Lippens, S. Ricard-Blum, X. Hanoulle, F. Penin, *Biochemistry* **2017**, *56*, 3029-3048.
2. X. Hanoulle, A. Badillo, J.-M. Wieruszeski, D. Verdegem, I. Landrieu, R. Bartenschlager, F. Penin, G. Lippens, *Journal of Biological Chemistry* **2009**, *284*, 13589-13601.
3. M. Dujardin, V. Madan, R. Montserret, P. Ahuja, I. Huvent, H. Launay, A. Leroy, R. Bartenschlager, F. Penin, G. Lippens, *Journal of Biological Chemistry* **2015**, *290*, 19104-19120.
4. R. L. Vold, *The Journal of Chemical Physics* **1972**, *56*, 3210-3216.
5. E. O. Stejskal, J. E. Tanner, *The Journal of Chemical Physics* **1965**, *42*, 288-292.
6. A. Chen, D. Wu, C. S. Johnson Jr, *Journal of the American Chemical Society* **1995**, *117*, 7965-7970.
7. F. Delaglio, S. Grzesiek, G. W. Vuister, G. Zhu, J. Pfeifer, A. Bax, *Journal of Biomolecular NMR* **1995**, *6*, 277-293.
8. J. J. Helmus, C. P. Jaroniec, *Journal of Biomolecular NMR* **2013**, *55*, 355-367.
9. M. Newville, T. Stensitzki, D. B. Allen, M. Rawlik, A. Ingargiola, A. Nelson, *Astrophysics Source Code Library* **2016**, ascl: 1606.1014.
10. W. Lee, M. Tonelli, J. L. Markley, *Bioinformatics* **2015**, *31*, 1325-1327.
11. C. D. Kroenke, J. P. Loria, L. K. Lee, M. Rance, A. G. Palmer, *Journal of the American Chemical Society* **1998**, *120*, 7905-7915.
12. N. Tjandra, A. Szabo, A. Bax, *Journal of the American Chemical Society* **1996**, *118*, 6986-6991.
13. A. Abragam, *The Principles of Nuclear Magnetism*, Oxford University Press, **1961**.
14. M. Lu, R. Ishima, T. Polenova, A. M. Gronenborn, *Journal of Biomolecular NMR* **2019**, *73*, 401-409.
15. M. Lu, S. Sarkar, M. Wang, J. Kraus, M. Fritz, C. M. Quinn, S. Bai, S. T. Holmes, C. Dybowski, G. P. Yap, J. Struppe, I. V. Sergeyev, W. Maas, A. M. Gronenborn, T Polenova, *The Journal of Physical Chemistry B* **2018**, *122*, 6148-6155.
16. H. M. McConnell, *The Journal of Chemical Physics* **1958**, *28*, 430-431.
17. G. Pagès, S. V. Dvinskikh, I. Furó, *Journal of Magnetic Resonance* **2013**, *234*, 35-43.
18. D. F. Hansen, P. Vallurupalli, P. Lundström, P. Neudecker, L. E. Kay, *Journal of the American Chemical Society* **2008**, *130*, 2667-2675.
19. D. Verdegem, A. Badillo, J.-M. Wieruszeski, I. Landrieu, A. Leroy, R. Bartenschlager, F. Penin, G. Lippens, X. Hanoulle, *Journal of Biological Chemistry* **2011**, *286*, 20441-20454.

### $R_{2,eff}$ Pulse Program

```
/*
| Pulse sequence to measure 19F R2 via CPMG
| including an anti-ringing sequence
*/

/*--------------------------------
;     Parameters to set
; -------------------------------*/
#include <Avance.incl>
#include <Grad.incl>
#include <Delay.incl>
/*--------------------------------
;        define loop counter
; -------------------------------*/
define list<loopcounter> ncyc_cp = <$VCLIST>
define delay Tau_cpmg
    "Tau_cpmg = d17 - p1"

"l2=0"      ; loopcounter for CPMG experiments in vclist
"l5=0"      ; counter for number of pi pulses in actual CPMG
"l6=0"      ; counter for number of pi pulses during compensation
"p2=p1*2"
"d11=30m"
"d13=4u"

1 ze
 3m st0

 if "Tau_cpmg < 125u"
{
  4u
  print "Tau_cpmg delay too short!"
  goto HaltAcqu
}

2 6m
3 6m
4 6m
```

```
/* ---------------------------------
;     calculation of delay
; --------------------------------*/

  "l5 = trunc(ncyc_cp[l2]+0.3)"
  4u pl1:f1
  30m

/*--------------------------------
; calculation number of
; heating compensation pulses
; for the next experiment
; -------------------------------*/
   "l6 = l7 - l5"
  if "l5 > 60"
{
  4u
  print " Number of ncyc(l5, CPMG) too large. ncyc < 61"
  goto HaltAcqu
}
  if "l6 > 60"
{
  4u
  print " Number of ncyc(l6, Heat Comp) too large. ncyc < 61"
  goto HaltAcqu
}

/* ---------------------------------
;     heating compensation
; --------------------------------*/

  if "l6 > 0"
  {

5    Tau_cpmg
     (p1*2 ph2):f1
     Tau_cpmg
  lo to 5 times l6

6    Tau_cpmg
```

```
     (p1*2 ph2):f1
     Tau_cpmg
  lo to 6 times l6
  }

100u

/* --------------------------------
;     add extra pi pulses
; to avoid starting magnetization
; --------------------------------*/
extra1,125u
      (p1*2 ph2):f1
      125u
      lo to extra1 times 20

/* --------------------------------
;     this is the start
; --------------------------------*/
7 d1
  20u UNBLKGRAD

  4u
  p16:gp1
  d16

  (p1 ph1):f1

/* --------------------------------
;     CPMG block
; --------------------------------*/

if "l5 > 0"
{
8  Tau_cpmg
   (p1*2 ph2):f1
   Tau_cpmg
lo to 8 times l5
}

if "l5 > 0"
```

```
{

9  Tau_cpmg
   (p1*2 ph2):f1
   Tau_cpmg
lo to 9 times l5
}

p1 ph3

4u
p16:gp2
d16

4u BLKGRAD

/* --------------------------------
;     anti-ringing
; --------------------------------*/

d13
(p1 ph1):f1
d13
(p1 ph4):f1
d13
(p1 ph5):f1

goscnp ph31

3m
iu2

d11 st
lo to 2 times nbl

ru2

3m ipp1 ipp2 ipp3 ipp4 ipp5 ipp31
lo to 3 times ns

d11 wr #0    ; write the data
```

```
3m rpp1 rpp2 rpp3 rpp4 rpp5 rpp31
3m zd        ; clear the memory

lo to 4 times td0

HaltAcqu, 1m
exit



ph1= 0 0 0 0 0 0 0 0 0 0 0 0 0 0 0 0 2 2 2 2 2 2 2 2 2 2 2 2 2 2 2 2 1 1 1 1 1 1 1 1 1
1 1 1 1 1 1 1 3 3 3 3 3 3 3 3 3 3 3 3 3 3 3
ph2= 1 1 1 1 1 1 1 1 3 3 3 3 3 3 3 3 1 1 1 1 1 1 1 1 3 3 3 3 3 3 3 3 0 0 0 0 0 0 0 0 2
2 2 2 2 2 2 2 0 0 0 0 0 0 0 0 2 2 2 2 2 2 2 2
ph3= 2 2 2 2 2 2 2 2 2 2 2 2 2 2 2 2 0 0 0 0 0 0 0 0 0 0 0 0 0 0 0 0 3 3 3 3 3 3 3 3 3
3 3 3 3 3 3 3 1 1 1 1 1 1 1 1 1 1 1 1 1 1 1
ph4= 2 0 2 0 2 0 2 0 2 0 2 0 2 0 2 0 0 2 0 2 0 2 0 2 0 2 0 2 0 2 0 2 3 1 3 1 3 1 3 1 3
1 3 1 3 1 3 1 1 3 1 3 1 3 1 3 1 3 1 3 1 3 1 3
ph5= 0 0 2 2 1 1 3 3 0 0 2 2 1 1 3 3 2 2 0 0 3 3 1 1 2 2 0 0 3 3 1 1 1 1 3 3 2 2 0 0 1
1 3 3 2 2 0 0 3 3 1 1 0 0 2 2 3 3 1 1 0 0 2 2
ph31=0 2 2 0 1 3 3 1 0 2 2 0 1 3 3 1 2 0 0 2 3 1 1 3 2 0 0 2 3 1 1 3 1 3 3 1 2 0 0 2 1
3 3 1 2 0 0 2 3 1 1 3 0 2 2 0 3 1 1 3 0 2 2 0


;pl1 : f1 channel - power level for pulse (default)
;p1 : f1 channel -  90 degree high power pulse
;p2 : f1 channel - 180 degree high power pulse
;d1 : relaxation delay; 1-5 * T1
;d11: delay for disk I/O                  [30 msec]
;d17: CPMG delay tau [ tau - pi - tau]
;d20: fixed echo time
;l7:  Maximum of vclist
;vc : variable loop counter, taken from vc-list
;ns: 8 * n or 64*n
;ds: > 128
;td1: number of experiments = number of values in vc-list
;VCLIST: List with number of cycles ( tau - pi - tau)2
;d20: decreases probe ringdown even in the absence of aring
```
